# Ablation of colonic epithelial Reg4+ support cells induces Notch-independent regeneration and mesenchymal remodeling

**DOI:** 10.1101/2022.01.28.478243

**Authors:** Timothy W Wheeler, Anne E Zemper

## Abstract

The colonic epithelium harbors a complex network of adult stem cells that integrate signals from many supporting cells to assist in their decision making. In this study, we ablate an epithelial secretory support cell population characterized by Reg4 expression, to investigate the systemic impact on stemness-related cell signaling pathways. Ablation of these cells results in a hyperproliferative state as well as paradoxical activation of Notch signaling, with the proliferative effect continuing even during Notch inhibition. Reg4+ cell ablation also causes an unexpected remodeling of the mesenchyme. We observe increased presence of Pdgfra-high fibroblasts and an expanded network of smooth muscle myofibroblasts, suggesting that Reg4-ablation reorganizes signaling between epithelium and mesenchyme. These changes occur in the absence of any significant immunological inflammatory response. Our data demonstrate that Reg4+ cells are critical directors of homeostatic epithelial-mesenchymal signaling. Further, this ablation model is an *in vivo* system for probing cell-cell interactions in the colonic stem cell niche.

**Graphical Abstract:** 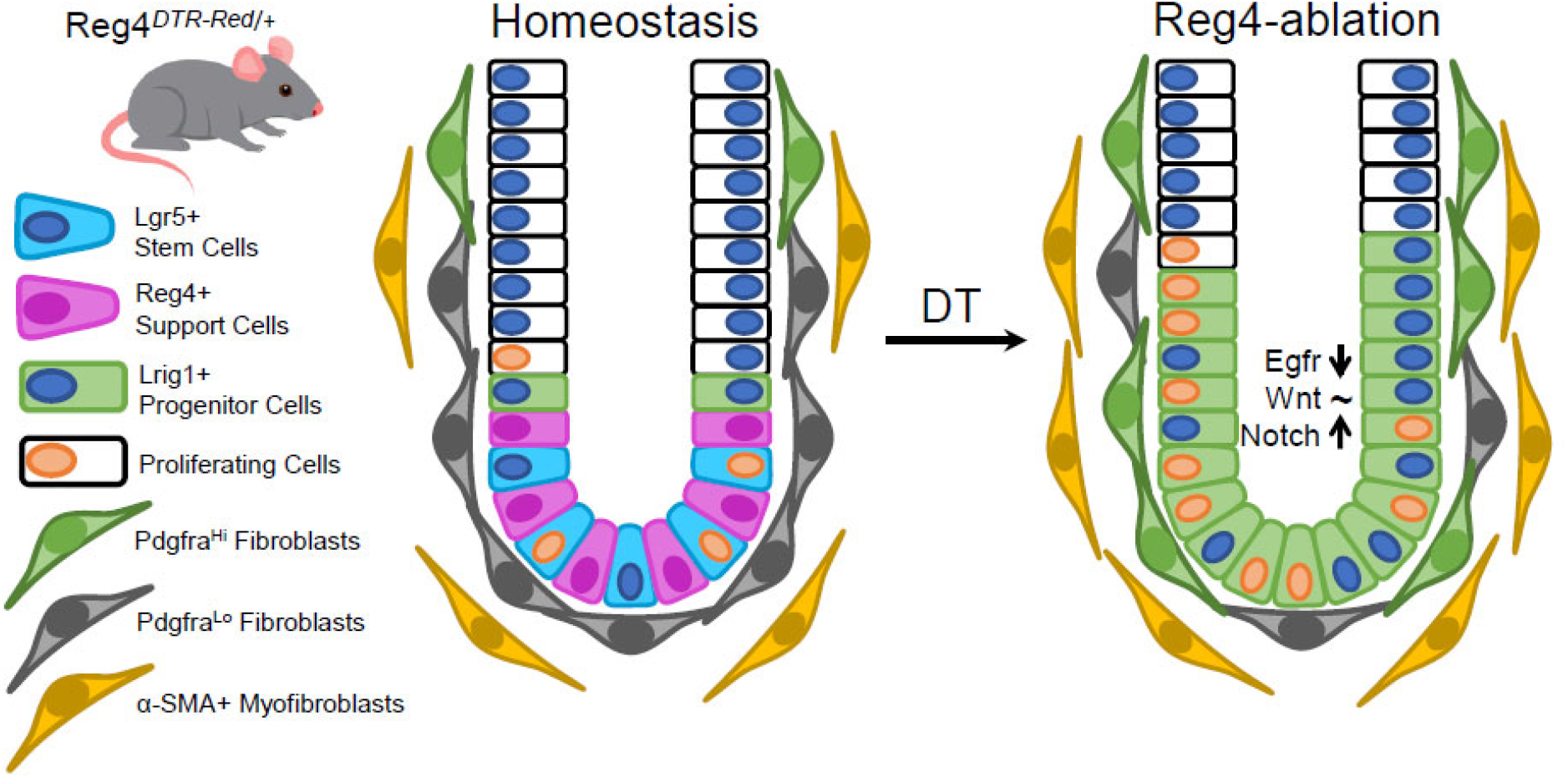

## Introduction

Epithelial stem cells are present across many genera in biology (Chacón-Martínez et al., 2018) and generally, stem cells act as population drivers - they give rise to more differentiated daughter cells which are responsible for the day-to-day duties of a given tissue. These cells often integrate extrinsic cues from both adjacent epithelial cells and their non-epithelial microenvironment to make decisions. The cells immediately surrounding the stem cell comprise its “niche”, and the stem cells exchange information with these cells throughout their life span (Chacón-Martínez et al., 2018; Meran et al., 2017). For tissues that constantly regenerate like the skin (Chacón-Martínez et al., 2018), blood (Wei & Frenette, 2018) and gastrointestinal epithelium (Chacón-Martínez et al., 2018; Meran et al., 2017), these cellular decisions are critical to maintaining the life of the tissue throughout an organism’s lifespan. Cells within these tissues have evolved mechanisms for handling changes within their environment and are generally thought of as dynamically-regulated and highly adaptable. Environmental changes can come in many forms. For the skin, it may be a burn or a scrape, resolved by epidermal stem cells dividing to generate additional skin cells to cover the wound, in combination with an inflammatory response that mitigates infection (Chen et al., 2018). But how are these injuries sensed? Why do some environmental changes elicit stem cell-based responses that differ from others? In short, what drives this adaptation to promote tissue maintenance over tissue death? To answer these questions, it’s helpful to take a reductionist approach by eliminating individual signals or cell populations, and then measuring the response in the stem cells. This is the approach we have taken to understanding colon adaptability.

In the colon, the cellular structure is critical to its function. The columnar epithelium is arranged as a single layer, acting as a barrier to intestinal contents, and is composed of millions of crypts (Potten, 1998). These crypts are U-shaped invaginations that harbor the stem cells, and their niche, at the crypt-base (Meran et al., 2017). The stem cells are highly proliferative, dividing once per day in mice and in humans, to generate a continual supply of differentiated daughter cells that are responsible for colon function. These stem cells have a unique molecular marker, Leucine Rich Repeat Containing G Protein-Coupled Receptor 5 (Lgr5), that distinguishes them from the other cells in surrounding epithelium (Barker et al., 2007). The cells that live on either side of these stem cells are differentiated cells and they also express a unique marker called Regenerating Islet-Derived Protein 4 (Reg4). Reg4+ cells express a myriad of ligands that bind to the stem cells to instruct a number of cellular functions, including regulation of proliferation and restricting differentiation. Recently, it was shown that ablation of these Reg4+ cells led to loss of Lgr5 expression in the remaining cells in the niche, and instead of collapsing, the crypts retained their unique structure and the cells within them continued to proliferate (Sasaki et al., 2016). The authors posited the cells remaining in the crypt may differentiate in response to ablation of Reg4+ cells, however this suggestion is at odds with the proliferative response they observed. Our study investigates this unexpected proliferative response, and we show that in the face of acute disruption the stem niche can immediately adapt to persist and recover. We show this is accomplished by a paradoxical shift in cellular signaling programs in the remaining cells of the crypt, accompanied by the reorganization of critical mesenchymal cell populations.

## Results

### Reg4 ablation induces transient hyperproliferation

Ablation of Reg4+ cells was previously shown to induce a loss of Lgr5+ stem cells without the loss of crypt-based proliferation (Sasaki et al., 2016). As stem cell loss generally induces a lack of proliferation and eventual organ atrophy, we first sought to more fully resolve this unexpected impact of Reg4-ablation on colon stem cells and proliferation throughout the course of this 6-day time period. We used *Reg4*^*dsRed-DTR/+*^ mice to ablate Reg4+ cells via daily diphtheria toxin (DT) injections as previously described (Sasaki et al., 2016), examining daily time points in a 6-day time course (Figure 1A). Control mice were injected with phosphate buffered saline (PBS) vehicle. As with the previous study, we observe the loss of Lgr5 expression by the 6th day of DT-driven Reg4+ cell ablation (Figure 1C, 1C’). We also observe an increase in total cell number per crypt, (Figure 1B), with the average cells per crypt cross-section increasing from 43 cells/crypt in vehicle to an average maximum of 53 at day 5 of Reg4-ablation. The driver of hyperplasia, cellular proliferation, increases soon after the start of time course (Figure 1D), with a significant increase in proliferation between 2-4 days of daily DT injections. The average maximum proliferation we observe is 33 Ki67+ cells/crypt at the 3-day timepoint, compared to an average of 11 Ki67+ cells/crypt in vehicle-injected controls (Figure 1D’). These data define a specific time course of hyperproliferation in response to ablation of Reg4+ cells within the first week of ablation.

**Figure 1.**
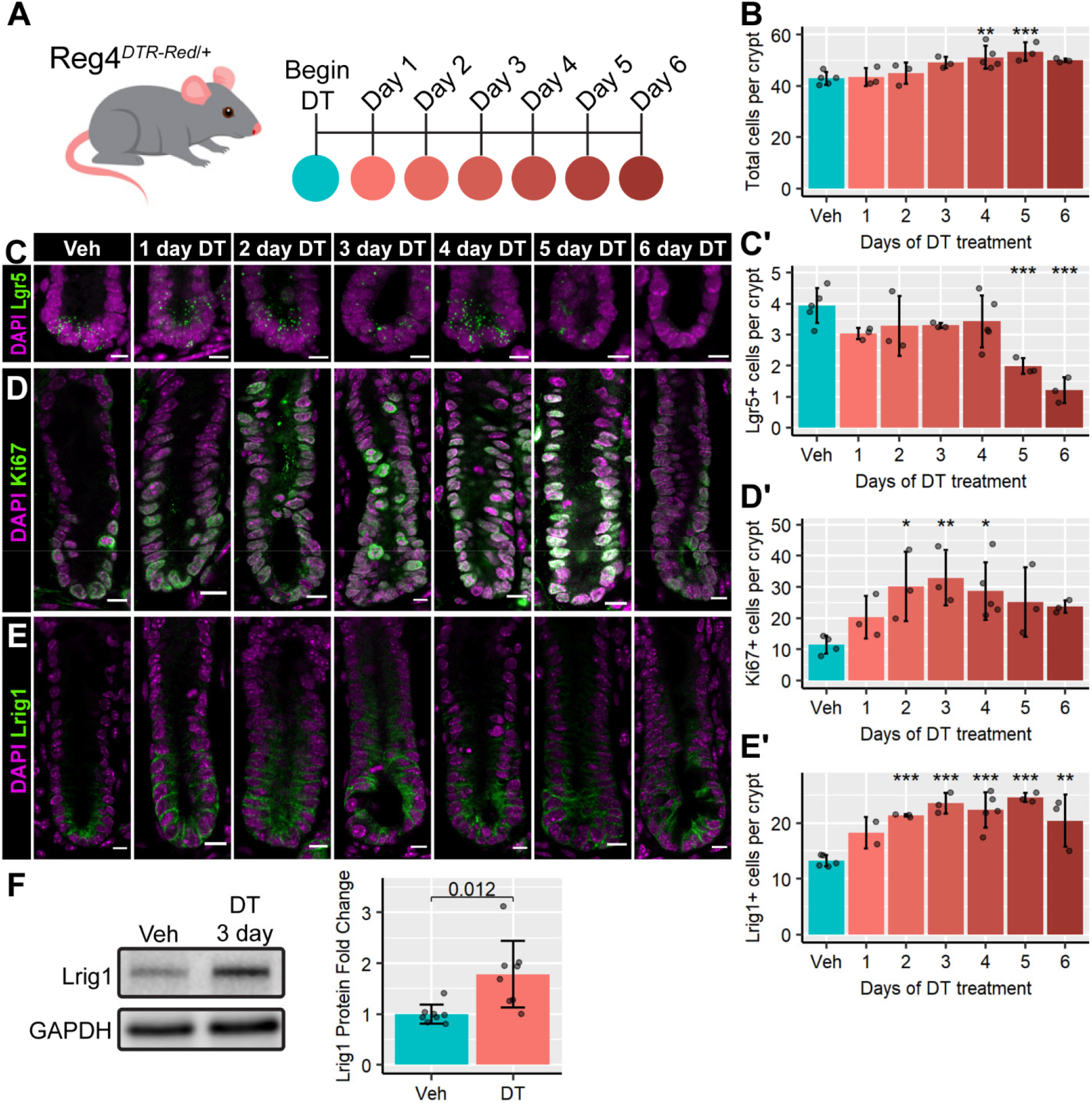
Reg4-ablation induces transient hyperproliferation. **(A)** Overview of Reg4-ablation time course, with diphtheria toxin (DT) or Vehicle (Veh; PBS) administered via intraperitoneal injection daily. Mice were sacrificed 24 hours after last injection, and colonic tissue collected 1-6 days after the start of injections. Vehicle injections were administered for 4 days. Diagram colors correspond to graph time points. **(B)** Total cells per well-formed crypt cross-section were measured across all stains (100-200 crypts per mouse). **(C)** *In-situ* hybridization for stem cell marker *Lgr5* (green) indicates a significant decrease at 5-6 days of DT treatment, quantified in **C’**. Images show maximum intensity projection of 5-μm z-stack. **(D)** Immunofluorescence for proliferation marker Ki67 (green) indicates a significant increase in proliferation increases significantly in response to Reg4-ablation, quantified in **D’**. **(E)** Immunofluorescence for progenitor marker Lrig1 (green) indicates a significantly expanded progenitor compartment during Reg4-ablation, quantified in **E’**. Nuclei are stained with DAPI (magenta). Graphs represent mean ± sd of mouse means, with individual mice represented as points. 15-25 crypts were measured per mouse for immunofluorescence and *in-situ* hybridization (n = 2-5 mice/group). Significance is calculated by nested Tukey test using crypts as random effect. *p*-values indicated: * < 0.05, ** < 0.01, *** < 0.001. All scale bars indicate 10 μm. **(F)** Western blot of purified mucosal protein indicates a 1.5-fold increase in Lrig1 in 3-day DT treated tissue compared to vehicle (n = 8 mice/group). GAPDH is used as a loading control. Significance indicated by *p-*value on graph, calculated by two-tailed unpaired Student’s t-test.

### Reg4 ablation expands a hyperproliferative progenitor cell pool

We sought the source of this transient hyperproliferation, by examining the progenitor epithelial cell pool to determine whether it is transiently expanding in response to Reg4-ablation, or whether an existing stem cell pool simply increases its proliferation rate. To address whether there were any dynamic, day-to-day changes to the pool of stem cells, we first examined the expression of *Lgr5*, which we observe is ultimately lost by day 6. We find no significant changes to the number of *Lgr5*+ cells until the 5-6 day time point when expression diminished (Figure 1C), which concurred with the previous study characterizing Reg4+ cells as support cells of the Lgr5+ cell population (Sasaki et al., 2016). We next examined the expression of Lrig1, an established stem and early progenitor cell marker in the colon (Powell et al., 2012). We see a significant increase in Lrig1-expressing (Lrig1+) cells and we observe this increased Lrig1+ population is maintained at an average of 1.7-fold above the vehicle-injected animals during this hyperproliferative phase (days 2-4) of Reg4-ablation (Figure 1E). This increase in Lrig1 expression was also verified by western blot, where Lrig1 is increased 1.5-fold at 3 days of Reg4-ablation (Figure 1F). To determine if Lrig1+ cells were responsible for the hyperproliferation we observe, we labeled proliferative cells within the crypt using a 2-hour pulse of 5-ethynyl-2’-deoxyuridine (EdU) on day 3 of Reg4-ablation (Figure 2A). These mice have a 5.3-fold increase in total EdU-labeled cells per crypt (Figure 2B), which agrees with the increase in Ki67 expression we observe (Figure 1D, D’). This expansion of EdU+ cells is concentrated within the Lrig1-expressing region of the crypt (Figure 2C). We quantified the EdU+;Lrig1+ cells and find the mitotic index of the Lrig1+ region increased from 0.12 in vehicle to 0.38 in the ablated crypts (Figure 2D). Taken together, our data suggest that crypt epithelial progenitor proliferation is significantly altered from homeostasis and the hyperproliferative response to Reg4+ cell ablation occurs in a dedicated pool of Lrig1+ cells.

**Figure 2.**
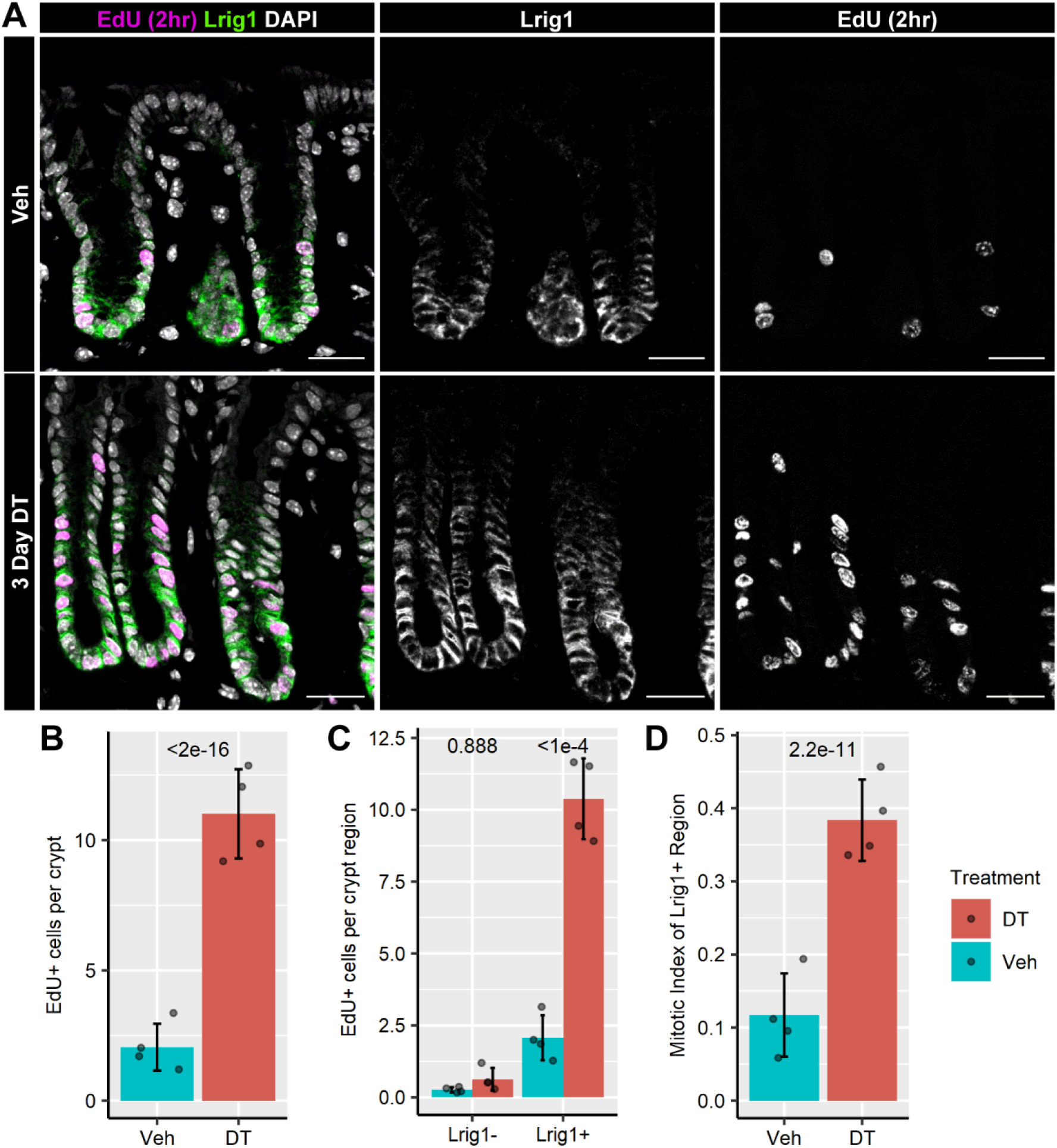
The hyperproliferation in Reg4-ablation is constrained to an expanded Lrig1-expressing region. **(A)** Colonic tissue from Reg4-DTR-Red/+ mice treated with Vehicle (Veh) or diphtheria toxin (DT) daily for 3 days, followed by a 2-hour trace of 5-ethynyl-2’-deoxyuridine (EdU) before mice were sacrificed. In the colored panels, representative images of DT and veh treated tissue were examined for the presence of proliferating cells (EdU labeling, magenta), nuclei by DAPI staining (white), and progenitor compartment by immunofluorescence for Lrig1 (green). Single color stains for Lrig1 and Edu are shown in the white-on-black panels. All scale bars indicate 25 μm. **(B)** Comparison of total EdU+ cells per crypt between Veh and DT. **(C)** EdU+ cells per crypt that do not coexpress Lrig1 (Lrig1-) or do coexpress Lrig1 (Lrig1+). **(D)**. Comparison of the fraction of total cells that proliferated (mitotic index) between Veh and DT in the Lrig1+ region. n = 4 mice/group, 40-60 crypts per mouse. Graphs represent mean ± sd of mouse means, with individual mice represented as points. Significance indicated by *p*-values on graphs, determined by nested Tukey Test using crypts as random effect.

### Reg4 ablation does not induce an inflammatory response

As epithelial hyperproliferation and hyperplasia are hallmarks of gut injury and repair (Davies et al., 2009; Zhang et al., 2012), we wondered whether this phenotype might be due to the stress of ablation inducing a macrophage-based inflammatory response that is also characteristic of injury models (Chen et al., 2018). To address this, we probed ablated tissues for expression of the macrophage marker F4/80, to determine if macrophage infiltration occurred after ablation (Figure 3A), as typical of inflammation after injury (Jones et al., 2018). In comparing vehicle-treated with DT-treated tissue, we see no significant difference from 1-4 days of treatment, using colons from mice subjected to dextran sodium sulfate (DSS) treatment as a positive control of macrophage infiltration (Novak et al., 2016). DSS-treated mice have a 2.3-fold increase in mean intensity of F4/80 compared to vehicle (Figure 3B). We do observe significant macrophage infiltration at 5-6 days after Reg4-ablation, but not until after the peak of hyperproliferation. Our results show that the expansion in proliferation in this model occurs in the absence of a classic macrophage-driven inflammatory response.

**Figure 3.**
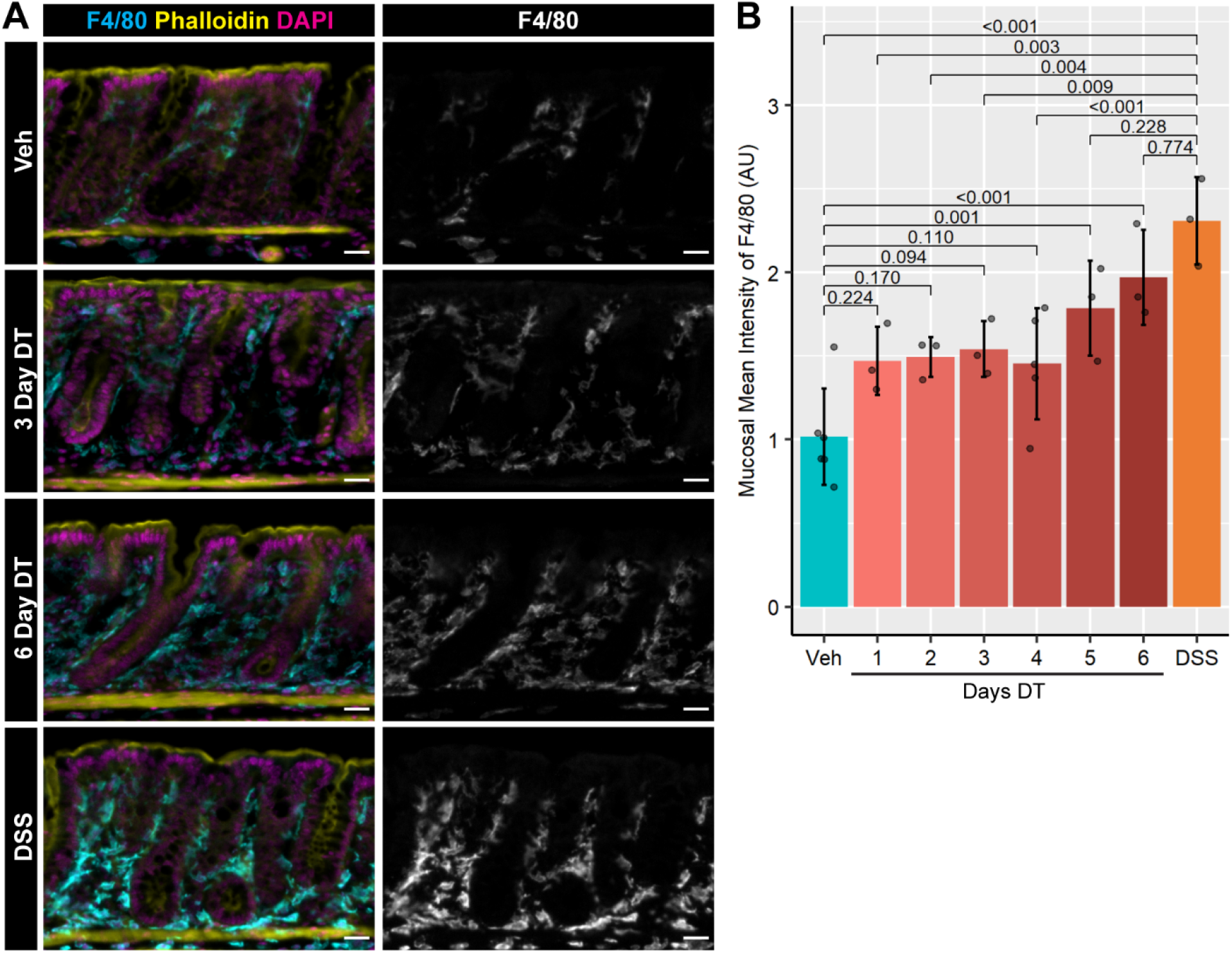
Macrophage-driven inflammation is not significantly increased in the initial proliferative response to Reg4-ablation. **(A)** Colonic tissue from Reg4-DTR/+ mice treated diphtheria toxin (DT), comparing to mice treated with vehicle (Veh) for 4 days as a negative control, and mice treated with 3% dextran sulfate sodium (DSS) in drinking water for 7 days followed by 3 days of recovery as a positive control for elevated macrophage infiltration. Presence of macrophages is indicated by F4/80 immunofluorescence (cyan), epithelial brush border is indicated by phalloidin immunofluorescence (yellow), and nuclei indicated by DAPI staining (magenta). Representative Images shown are from mice treated for 3 and 6 days with DT. Macrophage infiltration for all mice is in the single color (white on black) panels. **(B)** Quantification of mean intensity of F4/80 staining in total mucosal area across all groups indicates no significant difference between Vehicle and DT-treated mice, contrasted with significantly increased staining in DSS-treated tissue. Graphs represent mean ± sd of mouse means, with individual mice represented as points. n=3-5 mice/group, 5-7 images per mouse, with each image capturing 0.302 ± 0.060 mm^2^ mucosal area (mean ± sd). Significance indicated by *p*-values on graphs, determined by nested Tukey Test using images as random effect. All scale bars indicate 25 μm.

### Signaling pathways affecting stem cell maintenance are perturbed during Reg4 ablation

As epithelial hyperproliferation and hyperplasia are often driven by changes in specific cellular signaling programs, we next asked which signaling pathways were specifically perturbed in response to Reg4-ablation at the peak of proliferation (day 3). In the colon, there are numerous critical cell signaling pathways that guide stem cell decision making, and we prioritized three of these pathways in our analyses: Wnt (Nusse & Clevers, 2017), Epidermal growth factor receptor (Egfr) (Dubé et al., 2018), and Notch (Pellegrinet et al., 2011).

Wnt signaling is required for crypt-based proliferation (Nusse & Clevers, 2017), so we hypothesized that Wnt signaling would be increased immediately after Reg4-ablation due to the increase in the progenitor pool. An increase in variance of the Wnt signaling mediator, non-phospho-β-catenin (active), is detected in Reg4-ablated colonic tissue cell lysates (Figure 4A), suggesting perturbed Wnt pathway activation. Despite this, the downstream expression of Wnt targets remain relatively unchanged at this 3-day timepoint, with no significant changes to *Lgr5*+ cells (Figure 1C), *Lgr5* transcription (Figure S2), or cMyc patterning (Figure S3). As Reg4+ support cells provide Egfr and Notch support to the stem cells (Sasaki et al., 2016), we expected that these pathways would be significantly downregulated in the epithelium after Reg4-ablation. We find that Reg4-ablation causes a significant reduction in Egfr pathway activation (Figure 4A). While total Egfr expression is not significantly changed, phospho-Egfr expression (activated Egfr) decreases to 0.6-fold from expression levels in vehicle-treated mice. The downstream mediators of Egfr signaling, Erk1/2, are also significantly less expressed and activated, with Erk1/2 expression reduced to 0.7-fold and phospho-Erk1/2 reduced to 0.5-fold that of vehicle. We probed for Notch signaling using Notch1 NTM (Notch transmembrane-intracellular fragment) and NICD (Notch intracellular domain) as indicators of activated Notch (Zhdanovskaya et al., 2021), and find that Reg4-ablation causes a significant increase in both (Figure 4A). These findings of continued Notch signaling are especially notable as Reg4-lineage cells are thought to be the primary expressors of Dll1 and Dll4 (Figure S1,S2), the two primary Notch ligands expressed in the colonic epithelium (Pellegrinet et al., 2011). In Reg4-ablation, the number of cells expressing Notch ligands Dll1 and Dll4 are significantly decreased (Figure 4B,C) across all timepoints. Despite the loss of Notch ligands, we show continued Notch activation based on transcription of the Notch-activated transcriptional regulator *Hes1* (Figure S2) (Liang et al., 2019). Taken together, our results suggest signaling pathways affecting stem cell maintenance are perturbed during Reg4 ablation.

**Figure 4.**
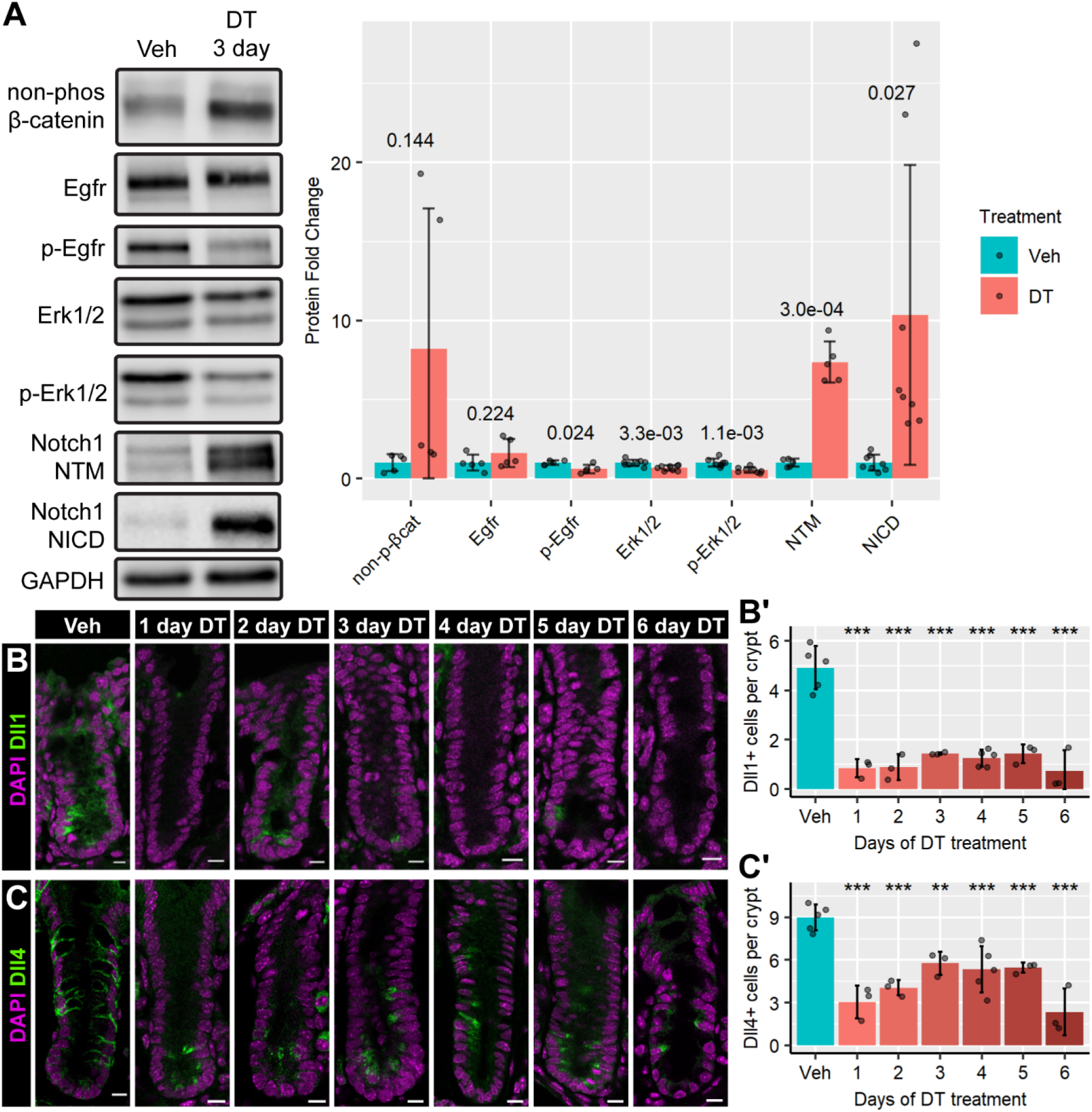
Reg4-ablation perturbs stemness-associated pathways. **(A)** Left panel: Western blot analysis comparing colonic mucosal tissue treated with Vehicle (Veh) or diphtheria toxin (DT) for 3 days. Egfr pathway proteins are detected with Egfr, p-Egfr, Erk1/2 and p-Erk1/2. Wnt signaling is detected by the presence of activated (non-phos) β-catenin, and Notch signaling was assessed by the detection of activated Notch1 (Notch1 transmembrane domain (NTM) and Notch1 intracellular domain (NICD)). Right panel: quantification of Western blot expression data (n=5-8 mice/group). GAPDH is used as a loading control. Significance indicated by *p-*values on graph, calculated by two-tailed unpaired Student’s t-test. **(B)** Immunofluorescence for Notch ligand Dll1 (green) indicates a reduction in Dll1-expressing crypt cells during Reg4-ablation time course, quantified in **B’**. **(C)** Immunofluorescence for Notch ligand Dll4 (green) indicates a reduction in Dll4-expressing crypt cells during Reg4-ablation time course, quantified in **C’**. Nuclear staining indicated by DAPI (magenta). Graphs represent mean ± sd of mouse means, with individual mice represented as points. 15-25 crypts were measured per mouse (n = 3-5 mice/group). Significance is calculated by nested Tukey test using crypts as random effect. *p*-values indicated: * < 0.05, ** < 0.01, *** < 0.001. All scale bars indicate 10 μm.

### Notch signaling is dispensable for the colon’s proliferative response to Reg4-ablation

Notch regulation is associated with proliferation in the intestinal tract (Bohin et al., 2020; Droy-Dupré et al., 2012; Pellegrinet et al., 2011), thus our next step was to test whether Notch activation was necessary for Reg4-ablation induced hyperproliferation. We conducted a dual time course of Reg4-ablation and Notch inhibition using the pan-Notch inhibitor Dibenzazepine (DBZ) (Bohin et al., 2020; Droy-Dupré et al., 2012) in a staggered time course (Figure 5A). We initiated Reg4-ablation 24 hours prior to Notch inhibition in order to initiate the proliferative trajectory. At 24 hours after ablation, we injected DT and DBZ for a subsequent 2 days to allow Notch inhibition to take effect. Two hours before sacrifice, we administered an EdU pulse to capture the proliferation occurring at that time. We observe intact gross mucosal and crypt morphology in all experimental conditions of this time course, although the distribution of visible secretory cells varied greatly by condition. DBZ-treated crypts show an increased density of secretory cells compared to vehicle, while DT-treated crypts show reduced secretory cell density irrespective of DBZ treatment (Figure 5B). Activated Notch1 is reduced in DBZ-treated mice, even with DT treatment (Figure 5C). We find that DBZ treatment alone does not cause a significant change to proliferation or body weight relative to vehicle (Figure 5D,E), and DT+DBZ treatment does not significantly increase proliferation relative to DT alone (Figure 5D), though body weight declines significantly faster in dual treatment compared to all other groups (Figure 5E). Despite the lack of significant change to total proliferation within crypts following Notch inhibition, we detect a significant shift in the location of proliferative cells, comparing Reg4-ablated mice with and without Notch inhibition. With respect to cell position from crypt base (position 0), proliferation shifts further up in crypts treated with DT+DBZ relative to DT alone (Figure 5D). Together, our results demonstrate Notch signaling is dispensable for hyperproliferation induced by Reg4-ablation, and inhibition of Notch signaling after Reg4-ablation causes proliferation to occur higher in the crypts.

**Figure 5.**
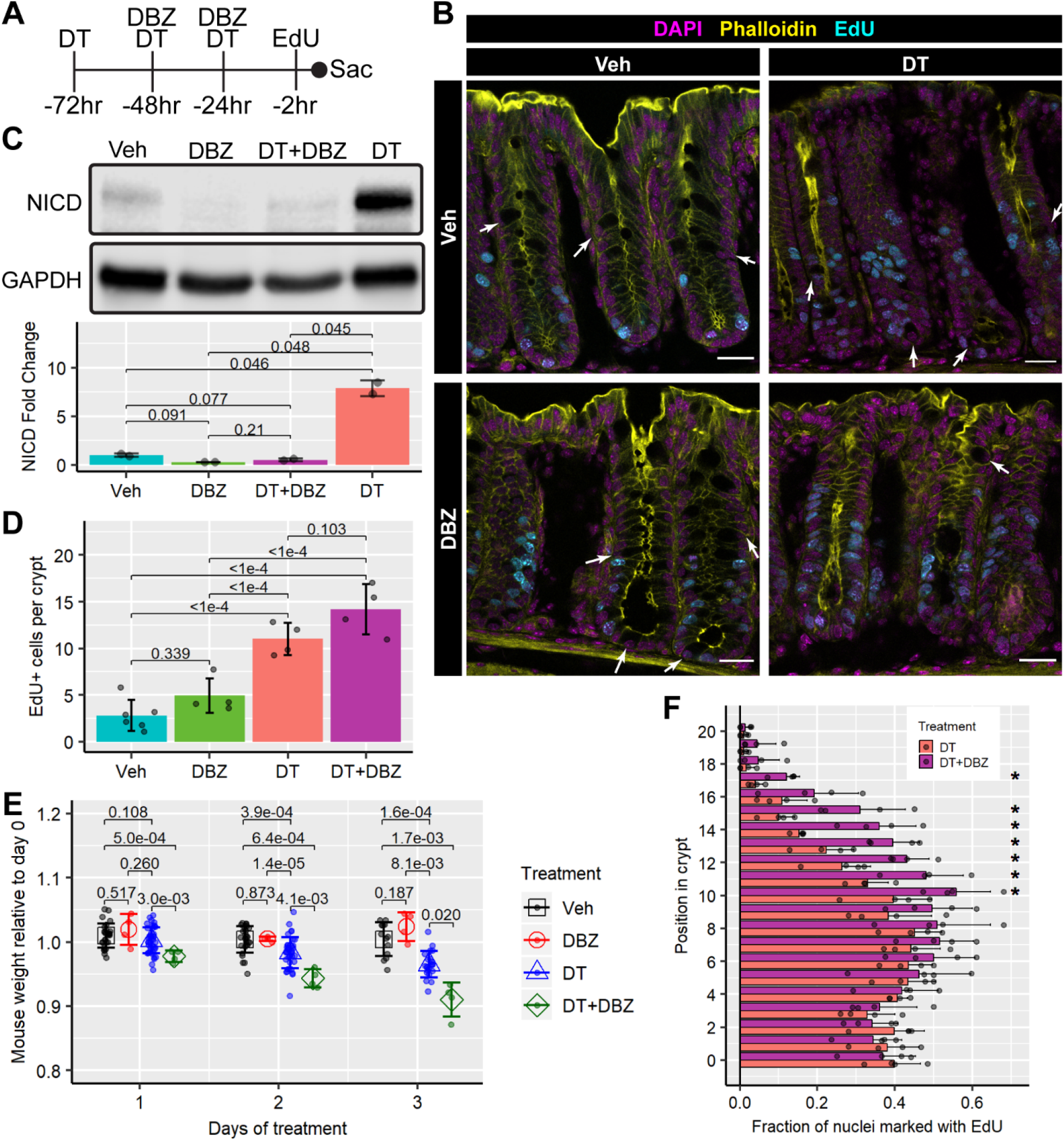
Notch inhibition does not downregulate hyperproliferation during Reg4-ablation. **(A)** Overview of dual Reg4-ablation and Notch inhibition time course to ablate Reg4-DTR-Red-expressing cells with diphtheria toxin (DT) and inhibit Notch signaling with *γ*-secretase inhibitor XX dibenzazepine (DBZ), followed by a 2-hour trace of 5-ethynyl-2’-deoxyuridine (EdU) before sacrifice. **(B)** Representative images of colonic tissue with EdU labeling (cyan) to detect proliferation, epithelial brush border by phalloidin immunofluorescence (yellow) and nuclei by DAPI staining (magenta). Arrows indicate examples of obvious secretory cells, visible as more bulbous cells highlighted by phalloidin. **(C)** Western blot of Notch1 intracellular domain fragment (NICD) shows successful inhibition of Notch by DBZ with and without DT, contrasting with Veh and DT (n=2 mice/group). GAPDH is used as a loading control. Significance indicated by *p-*value on graph, calculated by two-tailed unpaired Student’s t-test. **(D)** Comparison of total proliferation (EdU+) cells per crypt between all conditions, indicating significant differences between the DT treatment axis (n=4-6 mice/group). **(E)** A graph of mouse weights, relative to starting weight, over the time course. Precipitous weight loss occurs in DT+DBZ treatment compared to Veh and DT treatment alone. Veh and DT group data taken across all mice from study (n = 4 (DT, DT+DBZ) 28 (Veh) 43 (DT)). Significance comparisons indicated by *p-*values on graph, calculated by two-tailed unpaired Student’s t-test. **(F)** The fraction of proliferating cells per position relative to crypt base (position 0) is compared between DT and DT+DBZ treatments (n=4 mice/group). The proliferative region is shifted significantly higher on the crypt axis in DT+DBZ treated mice compared to DT. For immunofluorescence, 15-25 crypts measured per mouse. Graphs represent mean ± sd of mouse means, with individual mice represented as points. Significance indicated by *p*-values on graphs, determined by nested Tukey Test using crypts as random effect. All scale bars indicate 25 μm.

### The mesenchyme is remodeled during Reg4 ablation

Mesenchymal cells that surround the crypts are a critical component of colonic stem cell regulation (David et al., 2020; Degirmenci et al., 2018; Stzepourginski et al., 2017). We wondered whether mesenchymal cells, as a niche component, are also affected by Reg4-ablation. Groups of stem cell-‘supporting’ and - ‘antagonizing’ fibroblasts were recently characterized based on expression of Platelet-derived growth factor receptor-ɑ (Pdgfra)-low and -high expression, respectively. Pdgfra-high fibroblasts normally localize to the luminal surface of the colon in homeostasis (David et al., 2020). Using this marker, we examined patterns of Pdgfra expression relative to the crypt axis in ablated mice and find Pdgfra-high fibroblasts additionally localize to the crypt base (Figure 6A). To quantify this, we counted the number of epithelial cells, that are also adjacent to Pdgfra-high fibroblasts in the bottom half of crypts. Starting at Day 1 after ablation, the epithelial cells within this lower crypt compartment are 3 times as likely to be adjacent to Pdgfra-high fibroblasts, compared to vehicle-treated mice (Figure 6B). We also assayed the tissues for the expression of crypt-adjacent myofibroblasts that express α-smooth muscle actin (α-SMA) (Speca, 2012). We find that purified mucosal protein from Reg4-ablated tissue contains a 2.2-fold increase in α-SMA (Figure 6C). Comparing α-SMA and Pdgfra patterning to each other, we find that their expression patterns are distinct (Figure 6A). Our data show that mesenchymal cell relocation is cell-type dependent after Reg4-ablation. These patterns were consistent across all timepoints, suggesting that ablation of Reg4+ cells rapidly induces fibroblast remodeling within the adjacent mesenchyme.

**Figure 6.**
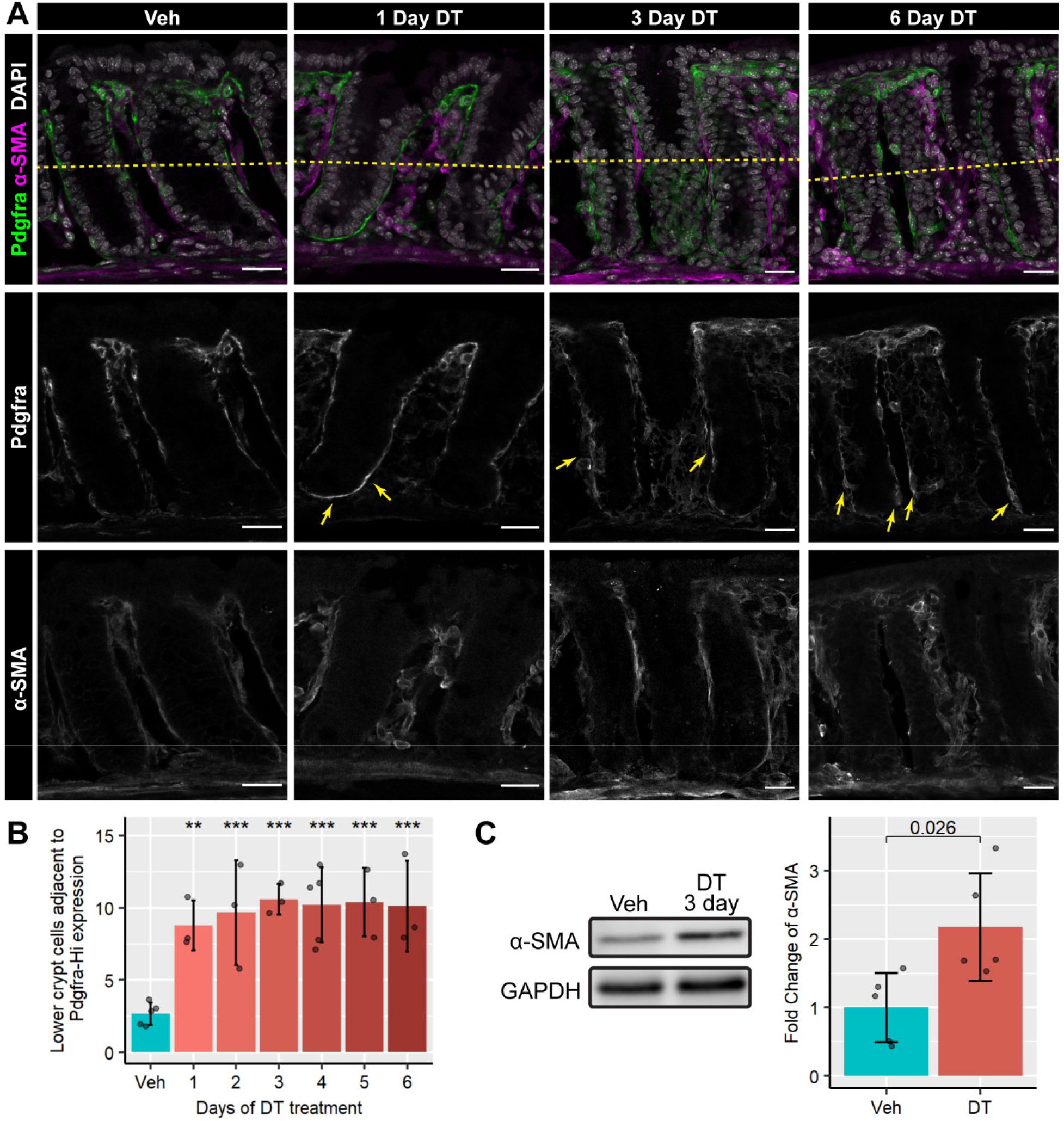
Reg4-ablation induces mesenchymal remodeling. **(A)** Colonic tissue from Reg4-DTR/+ mice treated diphtheria toxin (DT) daily for 1-6 days, compared to mice treated with vehicle (veh) for 4 days. Representative images show stromal fibroblasts indicated by Pdgfra immunofluorescence (green), structural myofibroblasts indicated by α-SMA immunofluorescence (magenta) and nuclei by DAPI staining (gray). Pdgfra-expressing stromal cells are detected as a distinct set of cells from α-SMA+ myofibroblasts. White-on-black images for Pdgfra and α-SMA are shown beneath the colored panel. In the single color panels, Pdgfra high-expressing cells (white) are apparent in DT-treated tissue around the crypt base (marked with arrows). α-SMA+ myofibroblasts (white) are present throughout the mucosa in all tissues. Scale bars indicate 25 μm. **(B)** Lower-crypt epithelial cells (below dotted line) adjacent to Pdgfra-high stromal cells are significantly increased at all time points of DT treatment. Graphs represent mean ± sd of mouse means, with individual mice represented as points. 15-25 crypts were measured per mouse (n = 3-5 mice/group). Significance is calculated by nested Tukey test using crypts as random effect. *p*-values indicated: * < 0.05, ** < 0.01, *** < 0.001. **(C)** Representative Western blot analysis of mucosal purified protein shows significant fold increase of α-SMA in mucosal isolate of 3-day DT treated tissue (quantification in right panel: n=5 mice/group). GAPDH is used as a loading control. Significance indicated by *p-*value on graph, calculated by two-tailed unpaired Student’s t-test.

## Discussion

In our study, we investigated induced ablation of Reg4+ secretory support cells and showed that in the face of acute disruption, the stem niche can immediately adapt to persist and recover. This recovery is achieved by induced epithelial hyperproliferation and expansion of an epithelial progenitor pool, and is accompanied by a paradoxical shift in cellular signaling programs in the remaining cells of the crypt and the reorganization of critical mesenchymal cell populations.

Reg4+ cells in the colon are often compared to the Paneth cells in the small intestine (van Es et al., 2019), as both are secretory support cells localized to the stem cell niche (Sasaki et al., 2016; van Es et al., 2019). Previous epithelial DTR ablation models in the gastrointestinal tract do not report a proliferative response to ablation (Castillo-Azofeifa et al., 2019; Tian et al., 2011; van Es et al., 2019), suggesting the response we observe is specific to colonic stem cells losing their epithelial support population. Our data suggests that Reg4+ cells are a distinct support cell population and have a distinct role that is different from that of the Paneth cell role in the small intestine. One open question our study does not address is whether the hyperproliferative response to Reg4-ablation is induced by loss of factors supplied by Reg4+ cells, or the act of ablating Reg4+ cells itself. Future studies resolving this question would provide useful insights into both the mechanisms underlying colonic epithelial homeostasis, as well as refine the applicability of this experimental model. We additionally find that *Spdef2/3* and *Muc2* are significantly downregulated by Reg4-ablation (Figure S2), and both are key components of the colon’s secretory lineage (Qin et al., 2021). This aids in characterizing the systemic impact of Reg4-ablation by illuminating the extent to which the broader secretory lineage is perturbed in this model. This is also contrasted with Paneth cell ablation in the small intestine, in which reserve secretory cells are readily able to compensate for the loss of Paneth cells (van Es et al., 2019); this notable difference in compensatory mechanism may explain some of the systemic response we observe and this represents an important area of future research.

We find that the Reg4-ablation model produces a transient hyperproliferative response reminiscent of other colonic injury models (Kiesler et al., 2015; Mizoguchi et al., 2020) yet it does this in the absence of many of the systemic effects that confound those models and prevent further analysis of molecular mechanisms that drive colonic homeostasis. By contrast, Reg4-ablated colons retain gross crypt and organ morphology, with no significant immune response during the acute period, which often follows tissue injury and obfuscates molecular analysis. Indeed, the Reg4-ablation model permits the dissection of the mechanisms underlying a proliferative response to injury, without other confounding factors present in existing models that induce systemic inflammation.

Disruption of the secretory lineage is associated with many other dysfunctions within gut injury models, and rescue of secretory cells improves the outcome after injury (Shinoda et al., 2010; Sun et al., 2018). These modes of injury are also linked to mesenchymal differentiation leading to fibrosis (Usunier et al., 2021), but it remains poorly understood how epithelial injury causes fibroblasts to promote fibrogenic activity. Our observations of altered fibroblast patterning in response to Reg4-ablation suggest that this system may be similar to other gut injury models in some respects. However, our data indicate that the Reg4-ablation is more of a “reductionist” model for epithelium-mediated colonic fibroblast activation, as the activation happens in absence of an inflammatory response. This model may aid in resolving less-understood mechanisms of fibroblast activation, compared to general injury models which have acute inflammation associated with them. Similar phenotypes are observed in colitis models, where extensive fibrosis is observed (Gregorio et al., 2017). These forms of fibrosis are linked to chronic bowel diseases (Speca, 2012), so a better understanding of the mechanisms through which fibroblast activation occurs may elucidate underlying pathways and help to develop strategies to mitigate fibrotic formation.

There are numerous cell signaling changes after Reg4-ablation. With regard to Wnt signaling, we observe fluctuations in active β-catenin after Reg4-ablation. As β-catenin is a structural protein, adherens junction remodeling that happens after Reg4-ablation may play a part in this phenomenon (Grainger & Willert, 2018). If it is an indicator of Wnt signaling perturbation, this change would likely come from surrounding fibroblasts that supply Wnt ligands (David et al., 2020), since epithelial cells do not express Wnt signaling ligands in the colon (Sasaki et al., 2016). Indeed, the changes we observe in β-catenin activation may be driven in part by the shift we detect in the fibroblasts after ablation. We observe Pdgfra-high fibroblasts shift to the crypt-base in response to Reg4-ablation, and these cells have previously been shown to express noncanonical Wnts and Bmp ligands (David et al., 2020). This shift in signaling would have a profound effect on epithelial cells if the shifted crypt base fibroblasts have similar expression to Pdgfra-high fibroblasts in homeostasis. Whether this shift in these mesenchymal cells directly causes the proliferative changes seen in the epithelial cells, represents a critical avenue for future research. To our knowledge, this is the first example of discrete, directed changes in the epithelial cell population causing a shift in mesenchymal cell localization. It will be important to define the mechanism for the translocation of these cells in future studies.

With respect to Notch signaling, we observe that Reg4-ablation induces a significant increase in Notch1 in the form of both NTM and NICD. Treatment with the γ-secretase inhibitor, DBZ, is sufficient to mitigate this increased Notch1 activation, suggesting that the increased Notch activation we see in Reg4-ablation is mediated through a canonical pathway. It remains unclear as to which process mediates this activation in spite of the loss of the majority of colon-specific Notch ligands, especially as we observe a paradoxical increase in activated Notch1. We noted a precipitous decline in body weight of Reg4-ablated mice treated with DBZ, suggesting that there are systemic redundancies between Reg4-expressing cells and Notch signaling within the mouse, although the colonic mucosa remains competent for proliferation. Within the colon, there was one notable change with respect to the localization of proliferation along the crypt axis. Normally the crypt base is the site of peak proliferation, as seen in homeostasis and Reg4-ablation. However, Notch-inhibited Reg4-ablated crypts have proliferation at the mid-crypt region and this may implicate Notch signaling as a component of crypt axis regulation. In summary, our data demonstrate the Reg4-ablation model as a novel model for the exploration of epithelial-mesenchymal signaling and Notch dysregulation.

## Materials and Methods

### Key Resources Table

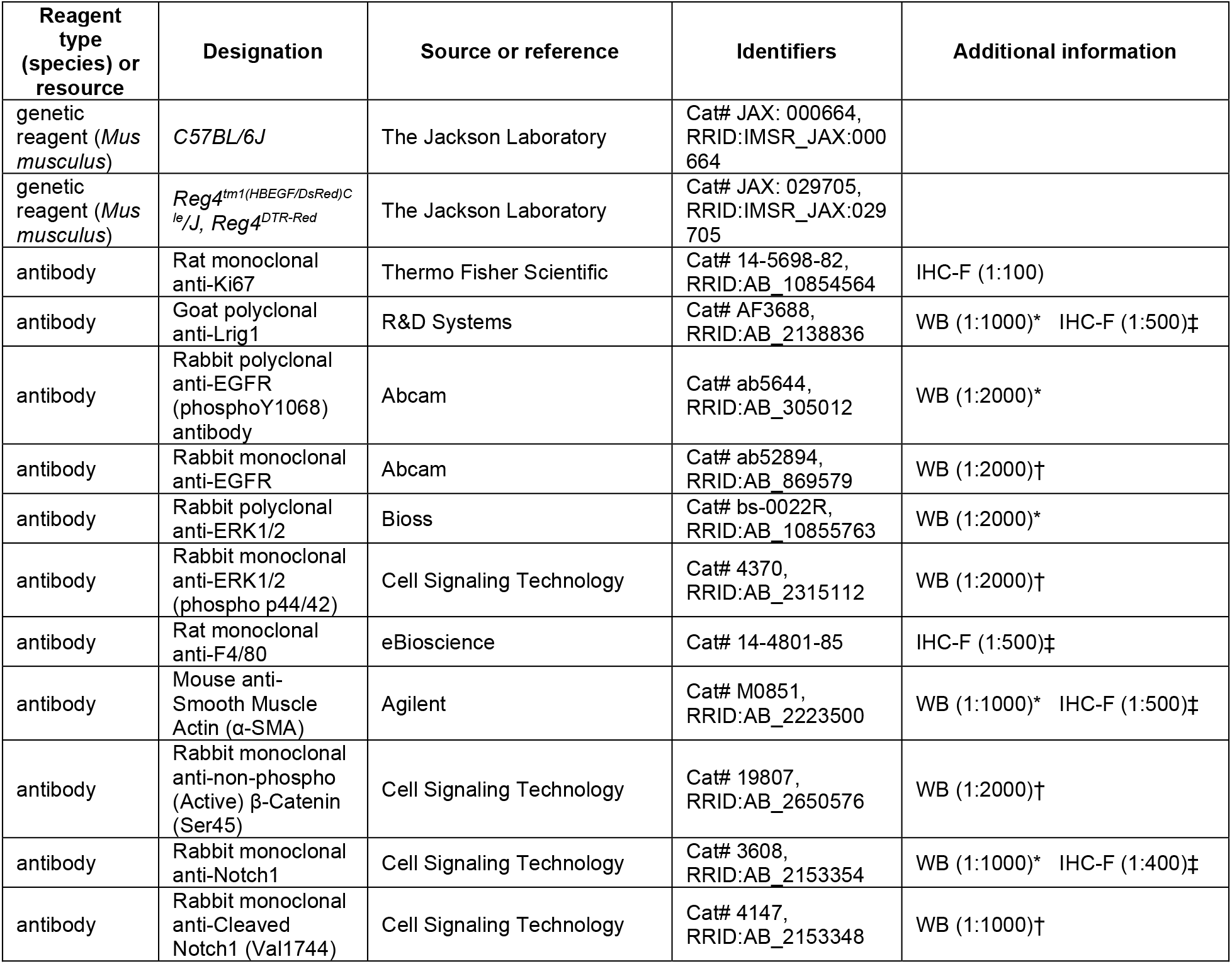
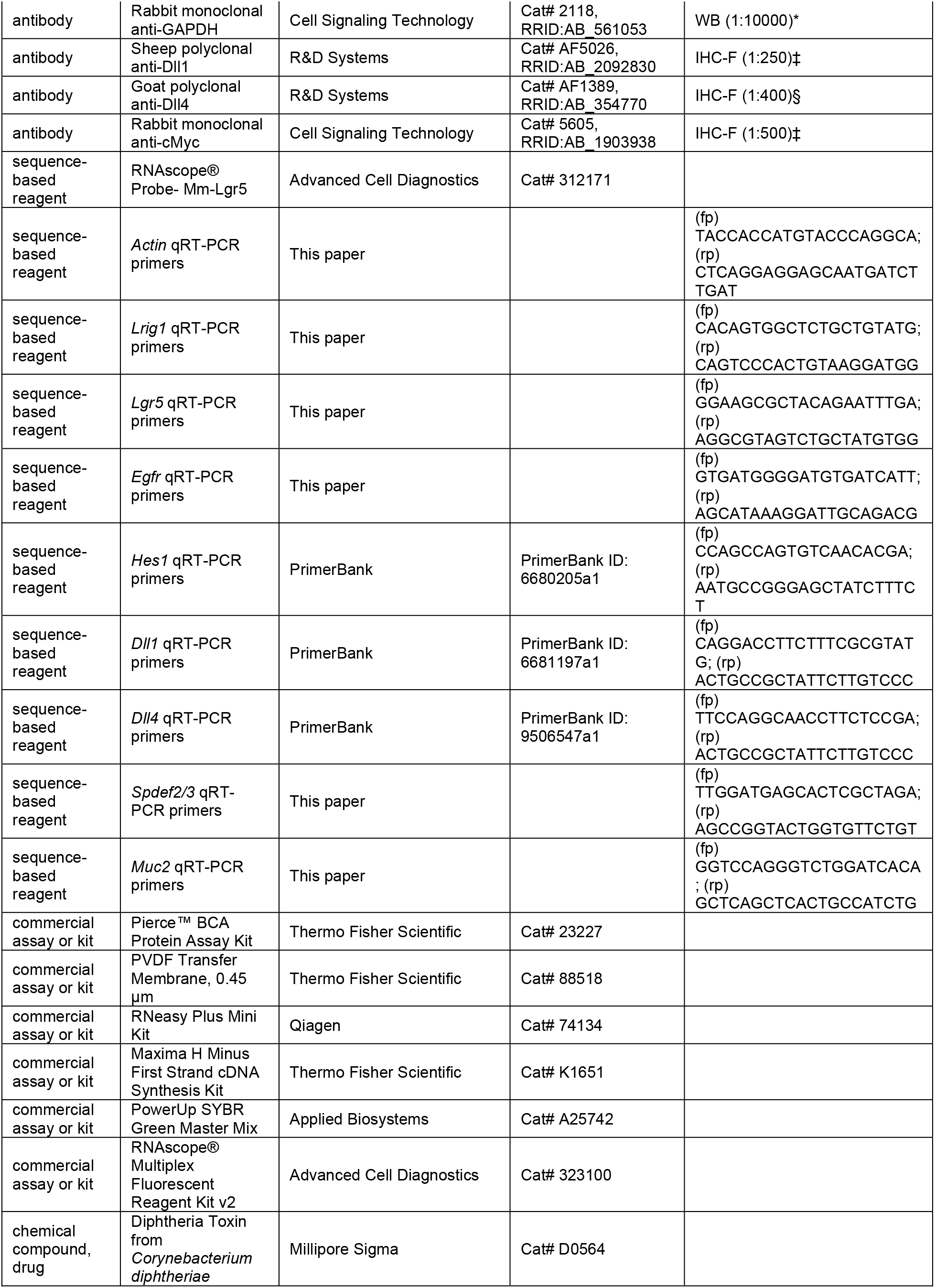
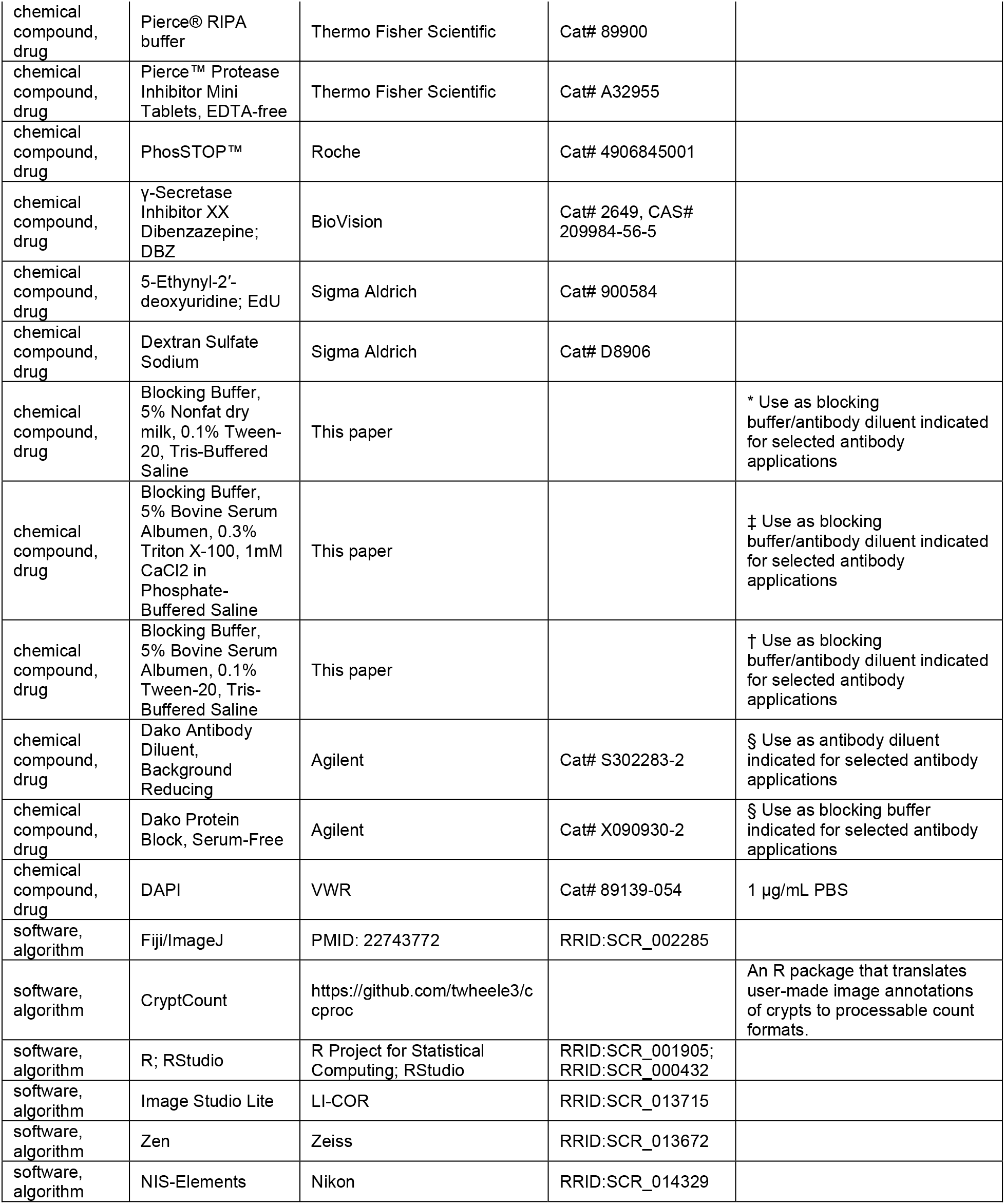

#### Mice

C57BL/6 (Jackson Laboratory (Jax), Bar Harbor, ME, USA) and Reg4-dsRed-DTR (Jax) mice were housed in a specific pathogen-free environment under strictly controlled light cycle conditions, fed a standard rodent lab chow and provided water ad libitum. Mice were sacrificed at 8-10 weeks of age by direct cervical dislocation. **Reg4-ablation**. Reg4-dsRed-DTR mice were administered Diphtheria toxin (CalBiochem, Billerica, MA, USA) at 50μg/kg body weight in 0.2mL PBS vehicle daily via intraperitoneal (IP) injection for 1-6 days of treatment, and were sacrificed 24 hours after last injection. Reg4-dsRed-DTR or C57BL/6 mice were administered 0.2mL PBS vehicle by IP injection for 4 days as a control group. **Notch inhibition**. Mice were administered γ-Secretase Inhibitor XX Dibenzazepine (DBZ) (BioVision #2649) at 10μmol/kg body weight (vehicle 0.5% (w/v) hydroxypropylmethyl cellulose, 0.1% (v/v) Tween-80, ddH_2_O) daily by IP injection for two days prior to sacrifice. **5-ethynyl-2’-deoxyuridine (EdU) tracing**. Mice were treated with 2mg EdU (Sigma-Aldrich, St. Louis, MO, USA) in 0.2mL DMSO/PBS vehicle by IP injection 2 hours prior to sacrifice. **Treatment with dextran sodium sulfate (DSS)**. DSS (Sigma-Aldrich) was dissolved in filtered drinking water and supplied ad libitum to C57Bl/6 mice for 7 days. Drinking water was switched to normal water on Day 8, then mice were sacrificed on Day 11 and colons were dissected for histology. All procedures were approved and performed in accordance with the policies of the University of Oregon Institutional Animal Care and Use Committee.

#### Histology

Colons dissected from mice were flushed with ice-cold PBS, flayed, and pinned flat, then fixed in 4% PFA/PBS for 60 minutes at room temperature with light oscillation. Fixed colons were washed 3×5 minutes in PBS and incubated in 30% sucrose overnight at 4°C, then blocked in OCT. 15 μm sections were taken on Superfrost™ Plus Slides (Fisher, Pittsburgh, PA, USA).

#### Immunohistochemistry

Slides were washed 3×3 minutes in PBS and blocked for 1 hour at room temperature in blocking buffer, then overnight at 4°C with primary antibody diluted in staining buffer. Slides were washed 3×3 minutes in PBS then stained with secondary antibody diluted in blocking butter. Slides were washed 3×3 minutes in PBS, counterstained with DAPI, and mounted with N-propyl gallate mounting medium. Slides were imaged via confocal microscopy using a Zeiss LSM-880 (Zeiss, Oberkochen, Germany) system for morphimetry, or a Nikon Eclipse/Ds-Ri2 (Nikon, Tokyo, Japan) for mean intensity analysis.

#### *In-situ* hybridization

Sections were labeled for *Lgr5* using RNAscope® Probe-Mm-Lgr5 (Advanced Cell Diagnostics (ACD), Newark, CA, USA) using RNAscope® Multiplex Fluorescent Reagent Kit v2 (ACD) as per manufacturer recommended protocol for fixed frozen sample (ACD TN 320535 Rev A, 323100-USM). In brief, slides were washed briefly in 1xPBS then boiled for 5 minutes in 1x Target Retrieval buffer, followed by two brief washes in ddH2O then once in 95% EtOH. Slides were dried then dammed with an ImmEdge hydrophobic barrier pen, then incubated with Protease III for 15 minutes at 40°C. Slides were washed briefly in ddH2O, then incubated at 40°C in sequence with Probe-Mm- Lgr5 (2 hours), AMP 1 (30 minutes), AMP 2 (30 minutes), AMP 3 (15 minutes), HRP-C1 (15 minutes), Opal-650 (1:2000 in TSA buffer) (30 minutes), with 2×2 minute washes with 1x Wash Buffer between hybridization steps. Slides were counterstained with DAPI and mounted with N-propyl gallate mounting medium. Slides were imaged via confocal microscopy using a Zeiss LSM-880 system.

#### Mucosal Tissue Isolation

Samples of isolated mucosal tissue were prepared from mice treated with either DT or vehicle daily for 3 days. The distal half of the colon was dissected out, flushed, flayed, and cut into 1-cm pieces and placed in 1 mM EDTA/5 mM DTT/PBS to incubate for 40 minutes on ice with gentle oscillation. Supernatant was decanted and replaced with 30mM EDTA/PBS, and the samples were incubated for 8 minutes at 37C with gentle agitation every 2 minutes. Supernatant was decanted and replaced with ice-cold PBS, and mucosal tissue was dissociated from muscle tissue by hand agitation (shaking at approx. 120 beats per minute for 1 minute intervals) until there were no apparent changes in turbidity. Visible muscle tissue was removed and dissociated mucosal tissue was pelleted at 1000xRCF at 4°C.

#### Western Blot

Isolated mucosal tissue was digested with 300uL Pierce® RIPA buffer (ThermoFisher, Waltham, MA, USA) treated with Pierce™ Protease Inhibitor Mini Tablets, EDTA-free (Thermo) and PhosSTOP™ (Roche, Basel, Switzerland) by syringing repeatedly through a 22-ga needle. Suspension was then centrifuged at for 5 minutes at 5000xRCF, then 5 minutes at 14000xRCF to clarify supernatant. Protein content was measured by BCA assay (ThermoFisher) to load 25μg of protein per western blot lane. Western blots were run with freshly prepared 10% acrylamide gels at 125 V and transferred to 0.45μm-pore PVDF membranes (ThermoFisher) at 55 V for 18 hours on ice. Membranes were dried, washed briefly with TBST, then blocked with specified blocking buffer for 1hr RT and incubated overnight with primary antibody diluted in blocking buffer at 4°C with light oscillation. Membranes were washed 3×3 minutes in TBST, then incubated for 1 hour in HRP-conjugated secondary antibody diluted in blocking buffer. Membranes were washed 3×3 minutes in TBST, then incubated for 5 minutes with Cytiva Amersham™ ECL™ Prime Western Blotting Detection Reagent (Cytiva, Marlsborough, MA, USA) prior to imaging for chemifluorescence on a LI-COR Odyssey Fc Imaging System (LI-COR, Lincoln, NE, USA). Membranes were stripped and restained up to 3 times, washing 2×10 minutes with mild stripping buffer (1.5% glycine, 0.1% SDS, 1% Tween-20, pH 2.2 in PBS), then 2×10 minutes with PBS and 2×5 minutes with TBST before re-blocking. Antibody and blocking conditions are described in Key Resources Table. Protein fold change was calculated versus vehicle by batch correcting based on vehicle response between membranes, normalizing to GAPDH expression as a loading control, then normalizing to vehicle mean.

#### qRT-PCR

Total RNA was isolated from mucosal tissue pellets using RNeasy Plus Mini Kit (Qiagen, Hilden, Germany). First strand cDNA was synthesized using 2 μg of total RNA using Maxima H Minus Kit according to manufacturer instructions (ThermoFisher). qRT-PCR was performed using PowerUp SYBR Green Master Mix (Applied Biosystems, Waltham, MA, USA) according to the manufacturer’s recommendations, and run using a Bio-Rad CFX96 Real-Time System (Bio-Rad, Hercules, CA, USA). Each target was run with three technical replicates per sample. The relative target gene mRNA levels were normalized to actin expression. Primers used are listed in Key Rescources Table.

#### Statistical analysis

Statistical analysis was performed using R. Morphometric data of blinded fluorescent images were hand annotated for crypts in Fiji (Schindelin et al., 2012) and processed in R for crypt structure using lab-produced software CryptCount (https://github.com/twheele3/ccproc). **Biological replicates** consisted of individual mice per treatment condition (n = 2-5 for immunofluorescence and *in situ* hybridization, 5-8 for WB and qRT-PCR). 15-30 well-formed crypts (visible in cross-section from muscularis mucosa to lumen) were counted per marker per mouse, and tested by Tukey test using crypts as random effect. WB and qRT-PCR were analyzed by unpaired two-tailed Student’s t-test.

## Abbreviations

αSMA: alpha Smooth Muscle Actin
DBZ: γ-Secretase Inhibitor XX Dibenzazepin
DSS: dextran sodium sulfate
DT: diphtheria toxin
DTR: diphtheria toxin receptor
DTT: dithiothreitol
EDTA: ethylenediaminetetraacetic acid
EdU: 5-ethynyl-2’-deoxyuridine
FISH: fluorescence in situ hybridization
IEC: intestinal epithelial cell
NICD: Notch1 intracellular domain
NTM: Notch1 transmembrane domain
OCT: Optimal Cutting Temperature compound
PBS: phosphate-buffered saline solution
PFA: paraformaldehyde
qRT-PCR: quantitative real-time PCR
RT: room temperature
TBST: Tris-buffered saline with 0.1% Tween-20
Veh: vehicle

---

**Supplemental Figure 1.**
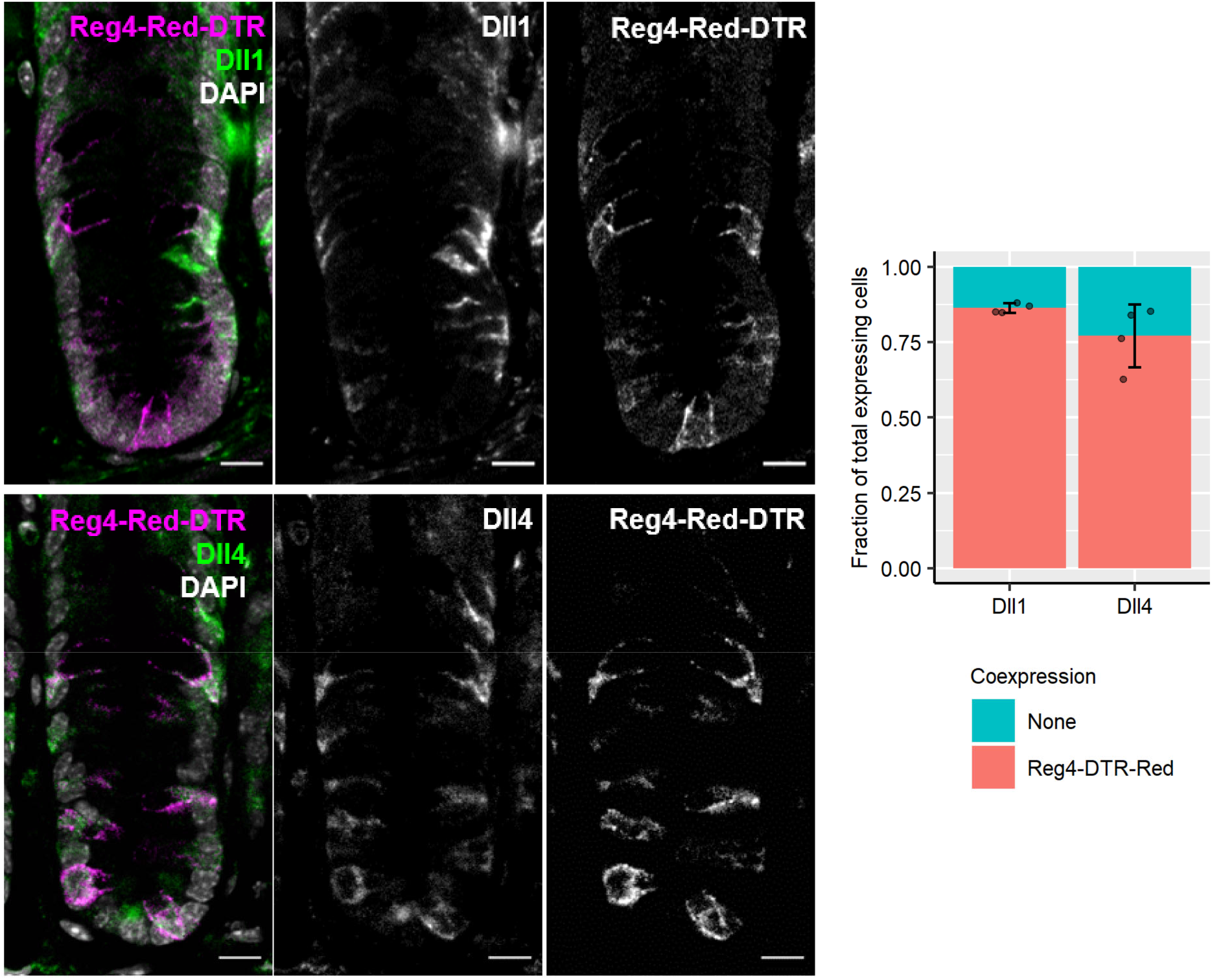
Notch ligand expression in the colonic crypt. Colonic tissue from Reg4-DTR-Red/+ mice. Notch ligands Dll1 and Dll4 are indicated by immunofluorescence (green), with Reg4-DTR-Red fluorescence indicated (magenta), and nuclei stained with DAPI (gray). Dll1 and Dll4 are primarily expressed by cells from a Reg4+ lineage (Reg4-DTR-Red) in the colonic epithelium, constituting 79% and 76% of total ligand-expressing cells, respectively (n=4 mice). Scale bars indicate 10 μm.

**Supplemental Figure 2.**
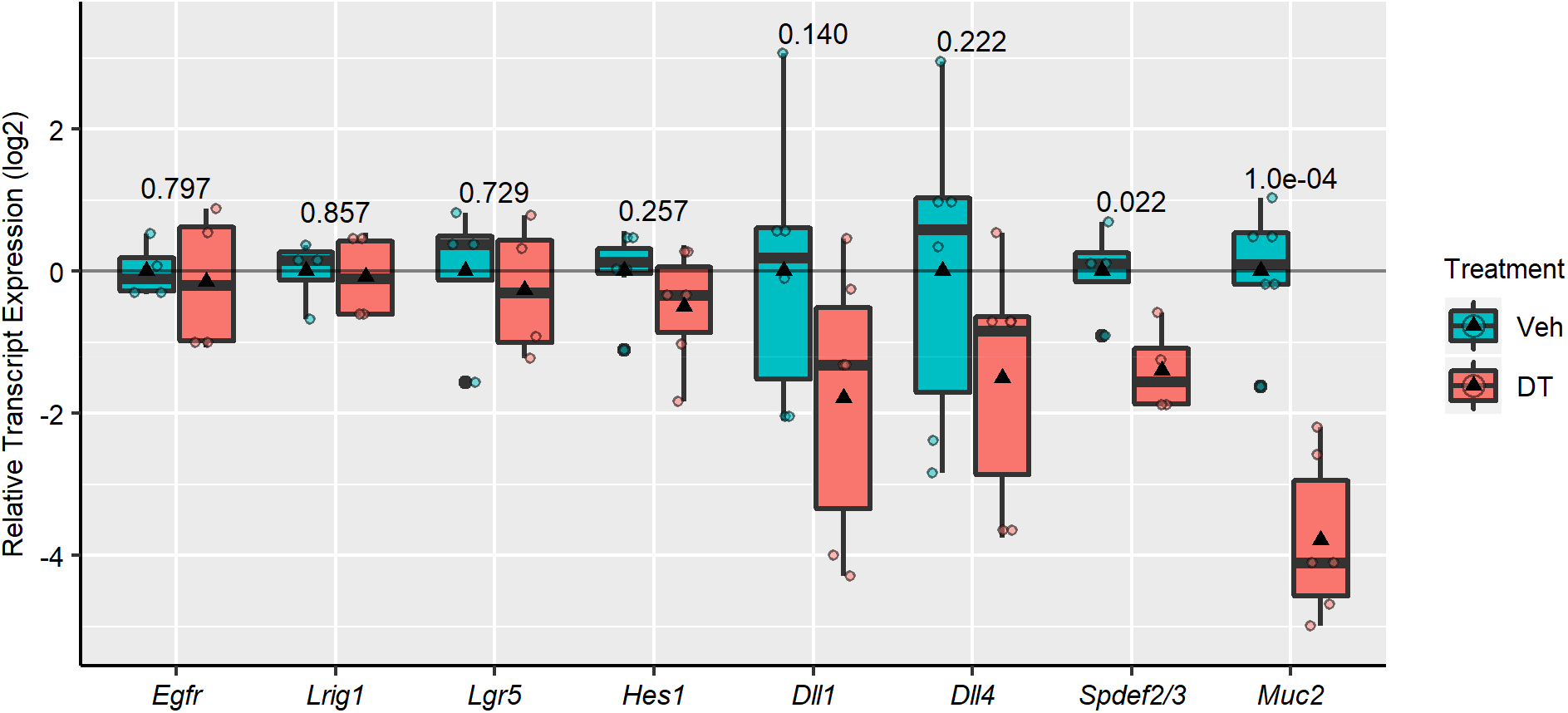
Reg4-ablation causes no significant change to relative transcription of several stemness-associated pathways, but reduces secretory transcription. qRT-PCR was performed on mucosal purified RNA from colonic tissue from Reg4-DTR-Red mice after 3 days of vehicle (veh) or diphtheria toxin (DT). No significant effect is observed in stemness-linked pathways (*Egfr*, *Lrig1*, *Lgr5*, *Hes1*), but relative transcription of secretory markers *Spdef2/3* and *Muc2* are significantly reduced. Boxplots represent quartiles with triangles as group mean (n=4-6 mice/group). Statistics taken with two-tailed unpaired Student’s t-test, *p*-values indicated on graph.

**Supplemental Figure 3.**
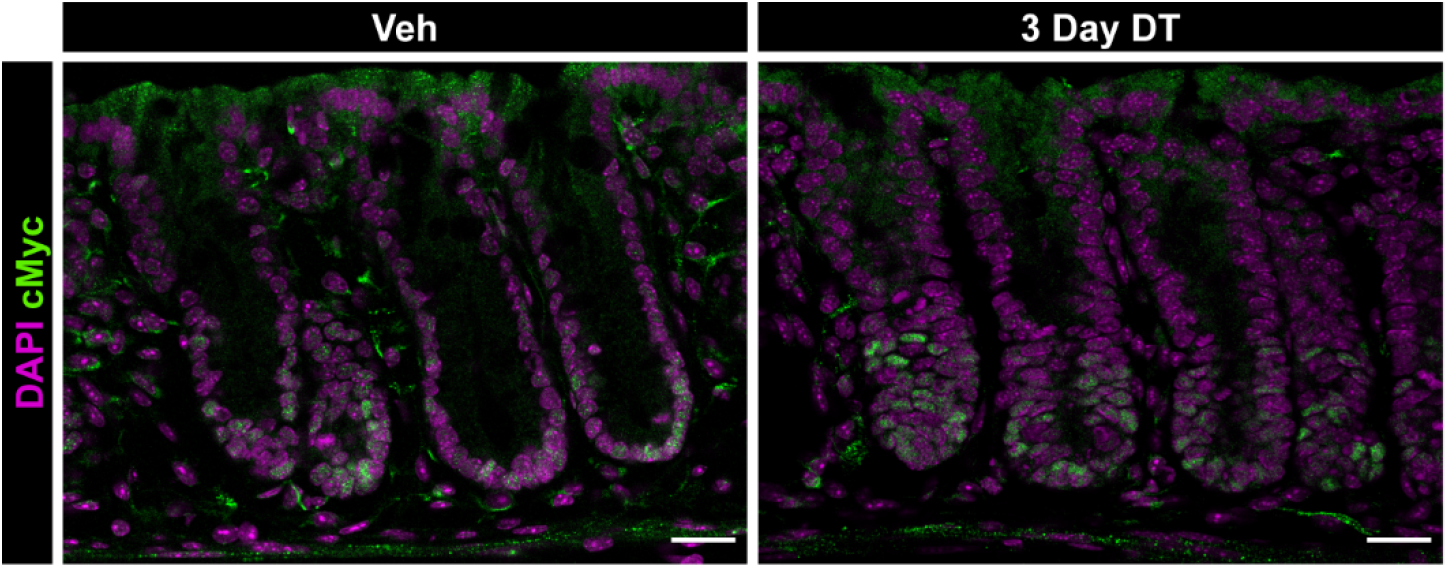
Patterning of cMyc, a downstream Wnt target gene, is not distinctly changed by Reg4-ablation. Colonic tissue from Reg4-DTR-Red/+ mice treated with vehicle (Veh) or diphtheria toxin (DT) for 3 days. Immunofluorescence of cMyc (green) shows no distinct changes in qualitative expression levels between Veh and DT. Nuclei are indicated by DAPI staining (magenta). Scale bars indicate 25 μm.

## Author Contributions

**Timothy W. Wheeler**

Contribution: Conceptualization, formal analysis, investigation, data curation, writing – original draft, visualization.

Competing interests: No competing interests declared. https://orcid.org/0000-0001-9518-4305

**Anne E. Zemper**

Contribution: Conceptualization, resources, writing – review and editing, supervision, funding acquisition.

Competing interests: No competing interests declared. https://orcid.org/0000-0001-8238-1406

## Acknowledgements

We thank Breanne Mohr for assistance with mouse colony management and experimental work. We gratefully acknowledge the use of the University of Oregon Genomics & Cell Characterization Core imaging facilities.

## Notes

### Competing Interest Statement

The authors have declared no competing interest.

